# Dendritic spikes in apical oblique dendrites of cortical layer 5 pyramidal neurons

**DOI:** 10.1101/2020.08.15.252080

**Authors:** Michael Lawrence G. Castañares, Greg J. Stuart, Vincent R. Daria

**Affiliations:** John Curtin School of Medical Research, Australian National University, 0200 ACT, Australia; Research School of Physics, Australian National University, 0200 ACT, Australia; Australian Research Council Centre of Excellence for Integrative Brain Function, Australian National University, 0200 ACT, Australia

**Author notes:** Corresponding author: Vincent Daria.

## Abstract

Dendritic spikes in layer 5 pyramidal neurons (L5PNs) play a major role in cortical computation. While dendritic spikes have been studied extensively in apical and basal dendrites of L5PNs, whether oblique dendrites, which ramify in the input layers of the cortex, also generate dendritic spikes is unknown. Here we report the existence of dendritic spikes in apical oblique dendrites of L5PNs. *In silico* investigations indicate that oblique branch spikes are triggered by brief, low-frequency action potential (AP) trains (~40 Hz) and are characterized by a fast sodium spike followed by activation of voltage-gated calcium channels. *In vitro* experiments confirmed the existence of oblique branch spikes in L5PNs during brief AP trains at frequencies of around 60 Hz. Oblique branch spikes offer new insights into branch-specific computation in L5PNs and may be critical for sensory processing in the input layers of the cortex.

## Introduction

Our capacity to decode how the brain works hinges in part on our understanding of how individual neurons process information. In the cortex of the mammalian brain, information processing is primarily performed by pyramidal neurons located in different layers of the cortical column. Of the different types of pyramidal cells in the cortex, the layer 5 pyramidal neuron (L5PN) plays a critical role in cortical processing. As its dendritic tree spans the entire cortical column, L5PNs receive and integrate information across all cortical layers (Spruston, 2008, Ramaswamy and Markram, 2015). For this reason, identifying the functional role of L5PN dendrites is crucial to decoding information processing in the cortex. While much is known about cortical processing in the apical tuft and basal dendrites of L5PNs (Nevian et al., 2007, Larkum et al., 2009), the apical oblique dendrites of L5PNs have received much less attention. Apical oblique dendrites of L5PNs receive almost a third of all synaptic connections from neighbouring L5PNs (Markram et al., 1997). In addition to these inter-cortical inputs, apical obliques of L5PNs are located in the primary input layer of the cortex (layer 4) and may receive direct input from the thalamus (Constantinople and Bruno, 2013). Hence, understanding how apical oblique dendrites process sensory information has important implications for cortical function.

An important feature that facilitates dendritic computation in L5PNs is the non-linear, regenerative activation of dendritic voltage-activated channels, which can lead to the generation of dendritic spikes (London and Hausser, 2005, Sjostrom et al., 2008, Major et al., 2013, Stuart and Spruston, 2015). While most studies on dendritic spikes in cortical pyramidal neurons have been performed in rodents, recent work indicates that human cortical pyramidal neurons can also generate dendritic spikes (Beaulieu-Laroche et al., 2018, Gidon et al., 2020). Previous works in cortical L5PNs indicates that dendritic spikes can be evoked in the basal (Kampa and Stuart, 2006, Nevian et al., 2007, Branco et al., 2010), apical trunk (Amitai et al., 1993, Kim and Connors, 1993, Stuart and Sakmann, 1994, Schiller et al., 1997, Stuart et al., 1997a, Larkum et al., 1999) and apical tuft dendrites (Larkum et al., 2007, Larkum et al., 2009, Harnett et al., 2015, Short et al., 2017, Beaulieu-Laroche et al., 2019, Fletcher and Williams, 2019). In contrast, whether dendritic spikes can also be evoked in apical oblique dendrites of L5PNs is largely unknown.

A number of studies suggest that oblique branches of L5PNs are endowed with active conductance that may support the generation of dendritic spikes. Large increases in intracellular calcium have been observed in oblique dendrites of L5PNs during epileptic discharges, suggesting that these dendrites express voltage-gated calcium channels (Schiller, 2002), while voltage imaging has revealed that back-propagating action potentials (bAPs) reliably invade the proximal oblique dendrites with minimal amplitude and time-course modulation (Antic, 2003). Interestingly, bAPs fail to invade some oblique branches, possibly due to heterogeneous expression of A-type potassium channels (Zhou et al., 2015). While these earlier studies suggest that the apical oblique dendrites of L5PNs are endowed with voltage-gated channels, direct evidence for locally generated dendritic spikes remains to be verified.

Here, we report dendritic spike generation in oblique branches of cortical L5PNs. Using the critical frequency protocol to evoke dendritic spikes with action potential (AP) trains (Larkum et al., 1999), we show in morphologically realistic models *in silico* and in real neurons *in vitro* that dendritic spikes are evoked in apical oblique dendrites of L5PNs at a critical frequency. In comparison to previous work where trains of four APs at around 100 Hz were required to evoke dendritic spikes at the nexus of the apical tuft (Larkum et al., 1999) and in basal dendrites (Kampa and Stuart, 2006) of L5PNs, we found that dendritic spikes in oblique branches are evoked by AP trains of just two APs at significantly lower frequencies (40 to 60 Hz). Oblique branch spikes offer new insights into branch-specific computations in L5PNs and may be critical for sensory processing in the input layers of the cortex.

## Results

Using a previously published multi-compartment model of a L5PN model (Shai et al., 2015), we investigated whether AP trains could evoke dendritic spikes in apical obliques dendrites, as has been previously shown to occur in the nexus of the apical tuft. We recorded the membrane potential at the soma as well as at the nexus of the apical tuft and also at various oblique dendritic branches (**Figure 1a**). Consistent with experimental findings (Larkum et al., 1999), a four-AP train at 100 Hz evoked a dendritic calcium spike at the apical nexus (**Figure 1b**). Surprisingly, we found that two-AP trains at much lower frequencies (36 Hz) evoked dendritic spikes in some oblique dendrites (**Figure 1c**). These oblique branch spikes had an approximate duration similar to that of dendritic spikes in the apical nexus. The generation of dendritic spikes at the apical nexus and in oblique dendrites led to a step-wise increase in the average membrane potential at the dendritic recording location (**Figure 1d**) as well as a stepwise increase in the ADP at the soma (**Figure 1e**) at the critical frequency.

**Figure 1.**
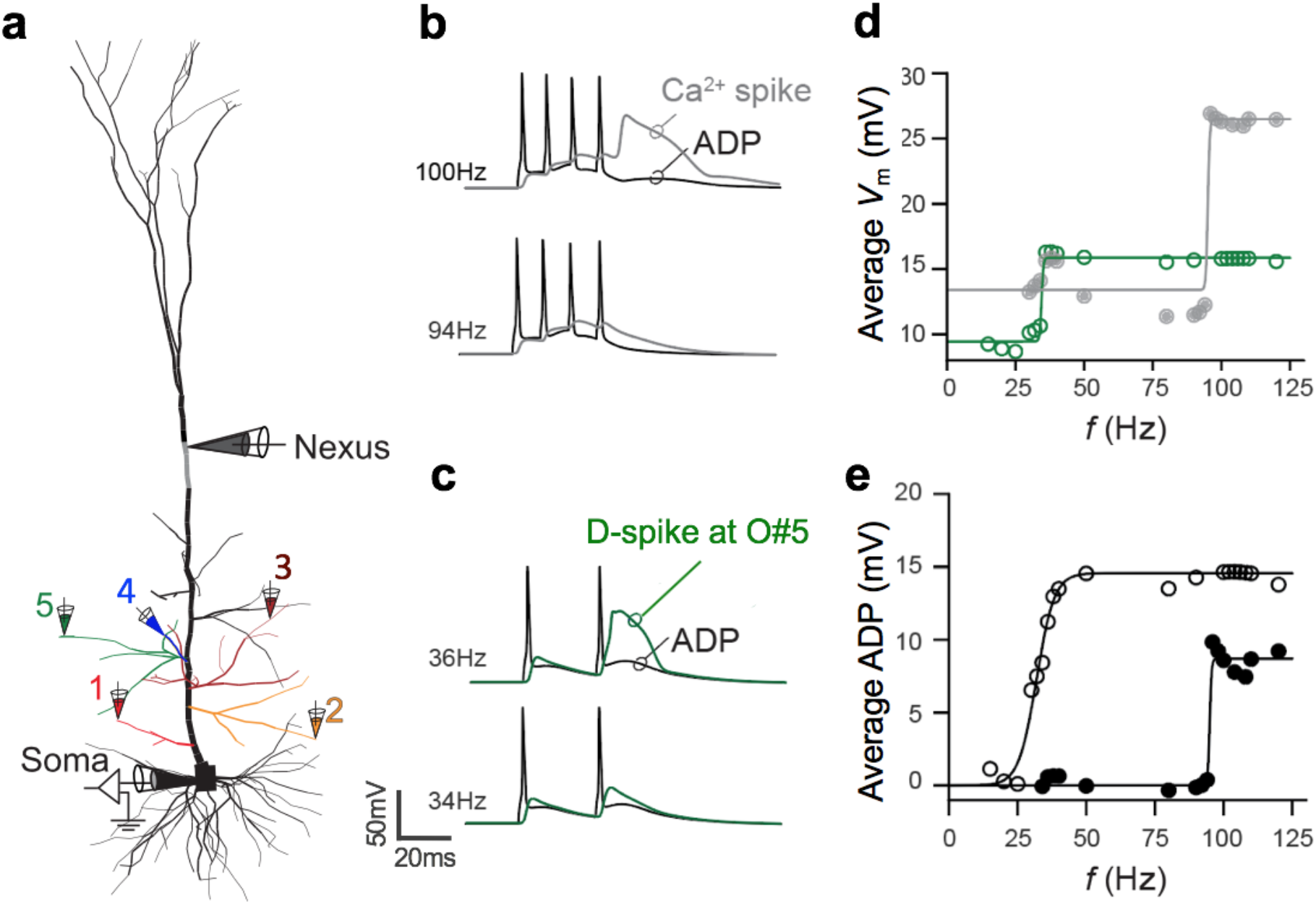
Dendritic spike generation at the obliques and nexus of the apical tufts. (**a**) The morphology of the L5PN multi-compartment model (Shai et al., 2015) with electrodes to indicate recording sites of the membrane potential at the soma, the apical nexus and oblique branches (numbered O#1 to O#5). (**b, c**) The membrane potential (*V*_m_) at the nexus (grey), and oblique dendrite O#5 (green) in response to four- and two-AP trains at the soma (black trace) at the indicated frequencies. (**d**) Average membrane potential (*V*_m_) over a fixed time interval (120 ms) at the apical nexus (grey) and oblique dendrite O#5 (green) during four-AP (filled circles) and two-AP trains (open circles) at different frequencies. (**e**) Average ADP voltage over a fixed time interval (5 ms) during four-AP (filled circles) and two-AP trains (open circles) at different frequencies.

We next evaluated the ionic mechanisms underlying oblique branch spikes by blocking voltage-activated sodium and low-voltage activated (LVA) and high-voltage activated (HVA) calcium channels in oblique dendrites in the model (**Figure 2a**). Blocking was performed by setting the channel densities to zero (i.e, 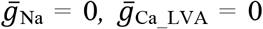, or 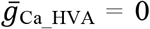) in specific oblique branches. Blocking LVA calcium channels did not affect the generation of oblique branch spikes (**Figure 2a**, 2^nd^ row). On the other hand, blocking HVA calcium channels abolished the broad ~20 ms depolarization in some obliques (e.g., O#2, O#3 and O#5), leaving only a fast spike of ~2 ms duration (**Figure 2a**, 3^rd^ row). Note that the dendritic spike in O#2 was already triggered with a single AP and was consequently abolished by blocking HVA calcium channels. Lastly, blocking voltage-activated sodium channels abolished oblique branch spikes (**Figure 2a**, 4^th^ row). From these numerical blocking experiments, we conclude that the oblique branch spike is generated by a fast sodium spike followed by a broad depolarization due primarily to activation of HVA calcium channels.

**Figure 2.**
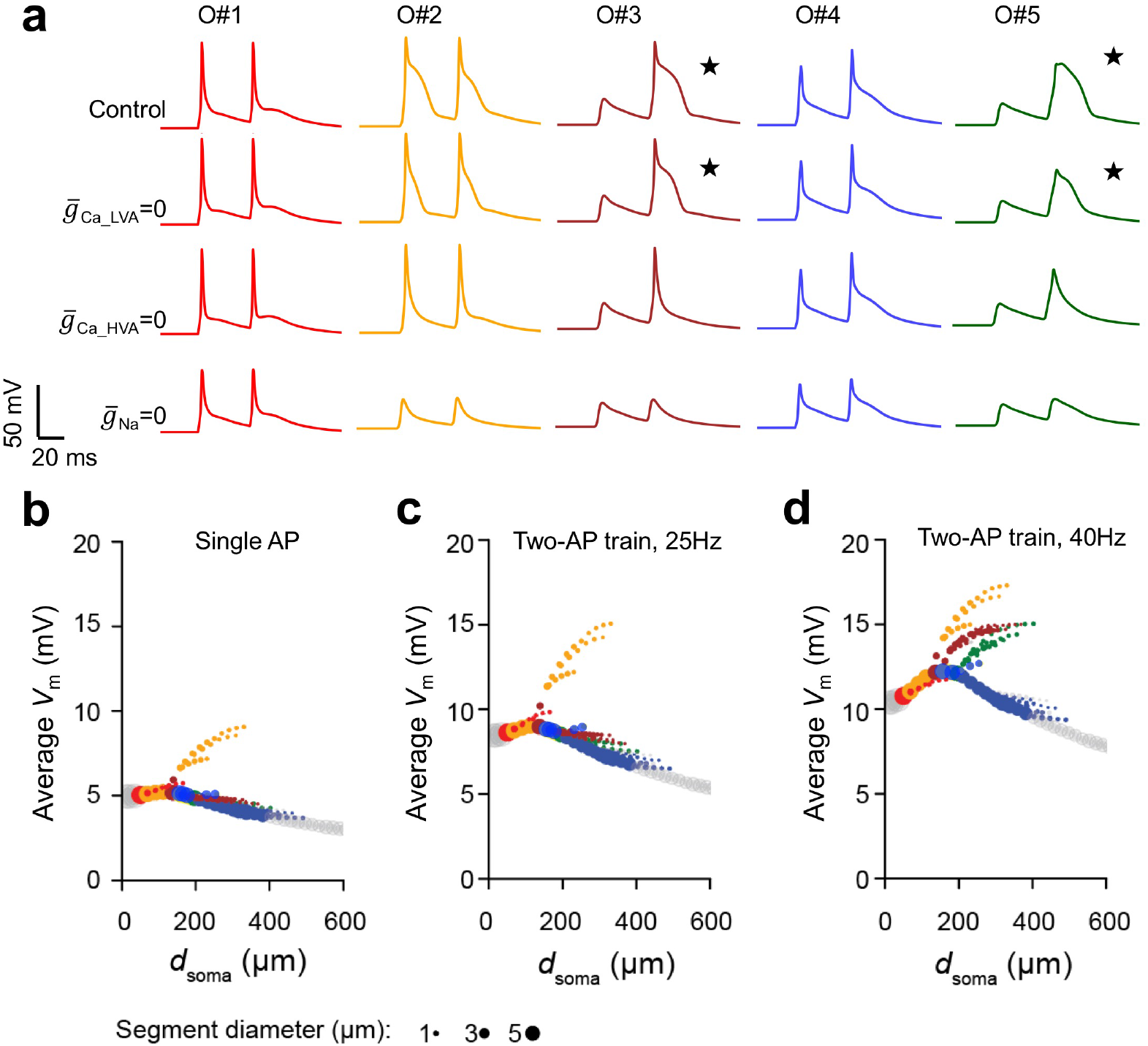
The excitability of the oblique branches in the L5PN model. (**a**) The membrane potential during a two-AP train at 36 Hz at oblique branches from O#1 to O#5 (see **Figure 1a**) in control (top) and after setting the 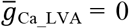 (second from top), 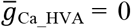 (second from bottom) and 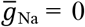 (bottom). Responses from branches that exhibited oblique branch spikes are indicated with ⋆. The average membrane potential over a constant time interval (120 ms) in oblique branches from O#1 to O#5 following: (**b**) a single AP, (**c**) a two-AP train at 25 Hz and (**d**) a two-AP train at 40 Hz. Colour code as in **a**.

The efficacy with which APs backpropagate into the dendritic tree is likely to influence the generation of oblique branch spikes. We therefore quantified the extent of AP backpropagation into apical oblique dendrites during single APs and two-AP trains by measuring the average membrane potential over a fixed time interval (120 ms) for each dendritic segment along apical oblique dendrites. All but one oblique dendrite (O#2) showed decremental backpropagation of single APs (**Figure 2b**). The same profile is observed during a two-AP train at 25 Hz with only O#2 showing regenerative backpropagation (**Figure 2c**). In contrast, increasing the frequency of the two-AP train to 40 Hz amplifies the average membrane potential along O#3 (brown) and O#5 (green), such that they now have a similar profile to O#2 (**Figure 2d**). The switch from decremental to regenerative backpropagation due to activation of voltage-gated ion channels further confirms the existence of frequencydependent dendritic spike generation in select oblique branches in this model.

Based on these modelling results, we tested experimentally whether oblique branch spikes could be evoked in L5PNs *in vitro*. We used two-photon multi-site calcium imaging to detect oblique branch spikes coupled with somatic whole-cell patch recording to evoke AP trains at defined frequencies. **Figure 3a** shows a schematic of a L5PN with 3 representative recording sites at the apical trunk (red) and two oblique branches (blue and green). Somatic current injection was used to evoke a single AP and a two-AP train, yielding dendritic calcium responses *C*_1_ and *C*_2_, respectively (**Figure 3b-d**). The calcium response during two-AP trains at frequencies up to 50 Hz was only slightly larger than that evoked during a single AP (*N* = 13 L5PNs) (**Figure 3b**). However, during two-AP trains at frequencies greater than 70 Hz, certain oblique dendrites exhibited calcium responses with significantly larger amplitude (**Figure 3c**; compare green with blue and red). A closer inspection at the temporal profile of these calcium transients indicated that a supra-linear increase in calcium influx occurs during the second AP (**Figure 3d**). The subtraction of the calcium response during a single AP from that observed during two-APs (Δ*C*_21_=*C*_2_-*C*_1_) showed a two-fold increase at certain obliques. Plots of Δ*C*_21_ for different frequencies of two-AP trains indicated a non-linear step increase in calcium occurred at frequencies around 70 Hz in some obliques (**Figure 3e**; green), but not in other oblique dendrites (blue) or the main apical trunk (red).

**Figure 3.**
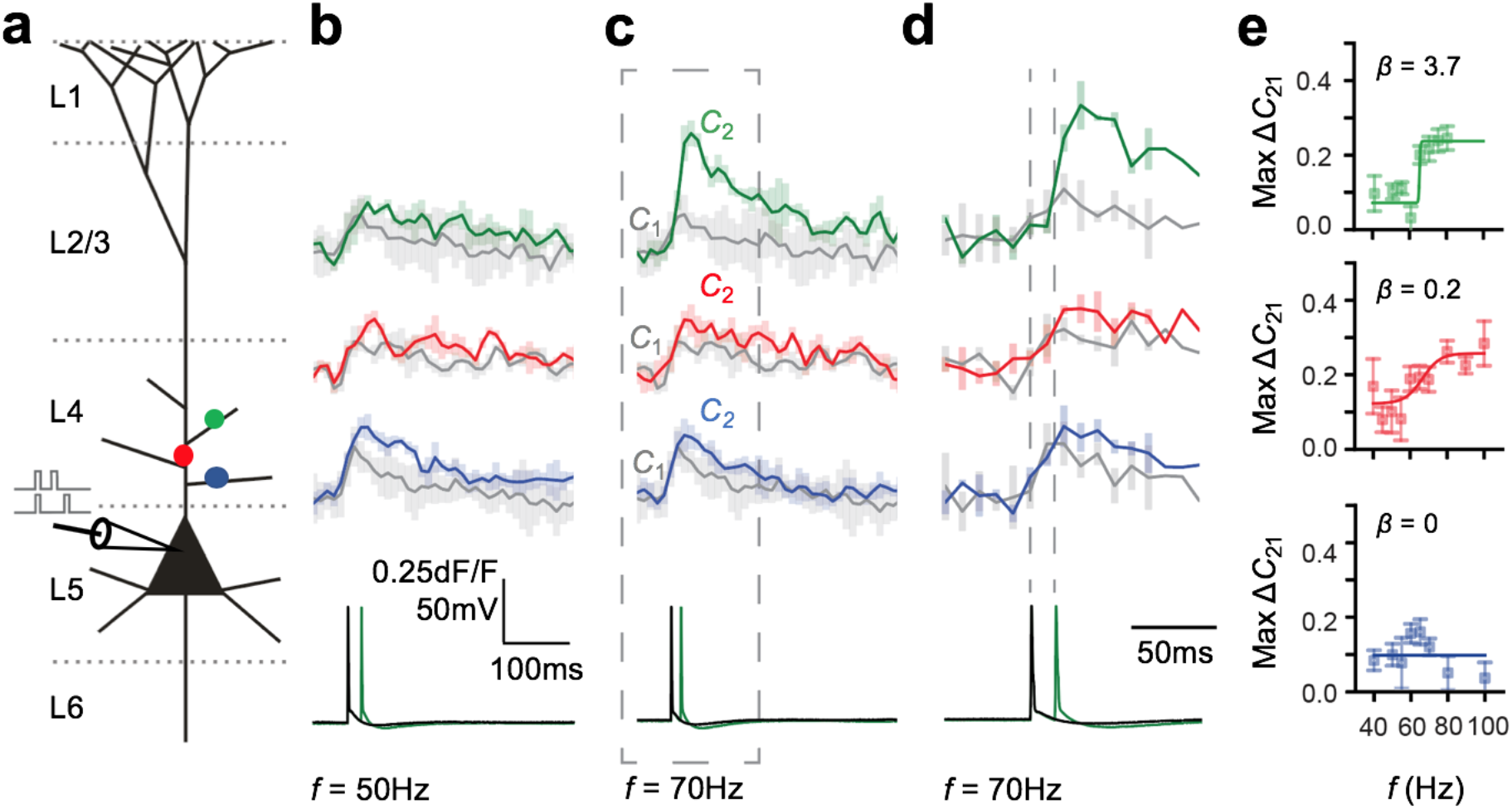
Calcium influx in different oblique branches following a two-AP train. (**a**) Schematic of a L5PN with representative recording sites at oblique branches (blue and green) and the apical trunk (red). (**b-c**) Calcium transients recorded at different dendritic locations (colour code as in **a**) during single APs (*C*_1_, grey) and two-AP trains (*C*_2_) at 50 Hz **(b)** and 70 Hz **(c)**. (**d**) Expanded response to a 70 Hz two-AP train (see box in **c)**. (**e**) Plots of Δ*C*_21_ at the three dendritic locations indicated in **a** during two-APs at different frequencies.

To determine if oblique branches generated an oblique branch spike or not, we went back to the model and analyzed the average membrane potential in different oblique branches during two-AP trains at different frequencies. Sigmoidal fits to plots of the average membrane potentials versus the frequency of AP trains allowed us to extract the slope (*β*_V_) and amplitude (*A*_V_) of this relationship. In the model, a threshold of *β*_V_ ≥ 0.3 and *A*_V_ ≥ 30 mV was sufficient to separate two types of responses (**Figure 4a** and **Supplementary Figure 3**). For experimentally measured calcium transients *in vitro*, we used a similar approach extracting the amplitude (*A*_C_) and slope (*β*_C_) of sigmoid fits to calcium transients during two-AP trains at different frequencies. We set *β*_C_ ≥ 0.3 as the common threshold that determines whether a dendritic spike was triggered in an oblique dendrite or not and set *A*_C_ ≥ 0.12 *dF/F*. This value captures 98% or more of the standard deviation of the noise across recorded calcium transients (*N* = 1794 traces from 11 L5PNs) and therefore sets a lower limit on our capacity to detect a change in calcium (**see Supplementary Figure 4**). We investigated a total of 38 oblique dendrites in 15 different L5PNs and found that these criteria effectively differentiated calcium-frequency responses associated with generation of dendritic oblique spikes from those that did not (**Supplementary Figure 5**). In total, six L5PNs showed oblique branch spikes in one or more oblique dendrites (**Figure 4b**). **Figure 4c** shows three representative 2P images of L5PNs annotating which oblique dendrites elicited oblique branch spikes (red dots). Consistent with the model, we found that only a small fraction of oblique branches in L5PNs elicited dendritic branch spikes. On average, in those obliques with branch spikes, the critical frequency for evoking oblique branch spikes during two-AP trains was 63±4 Hz (*N* = 6 L5PNs).

**Figure 4.**
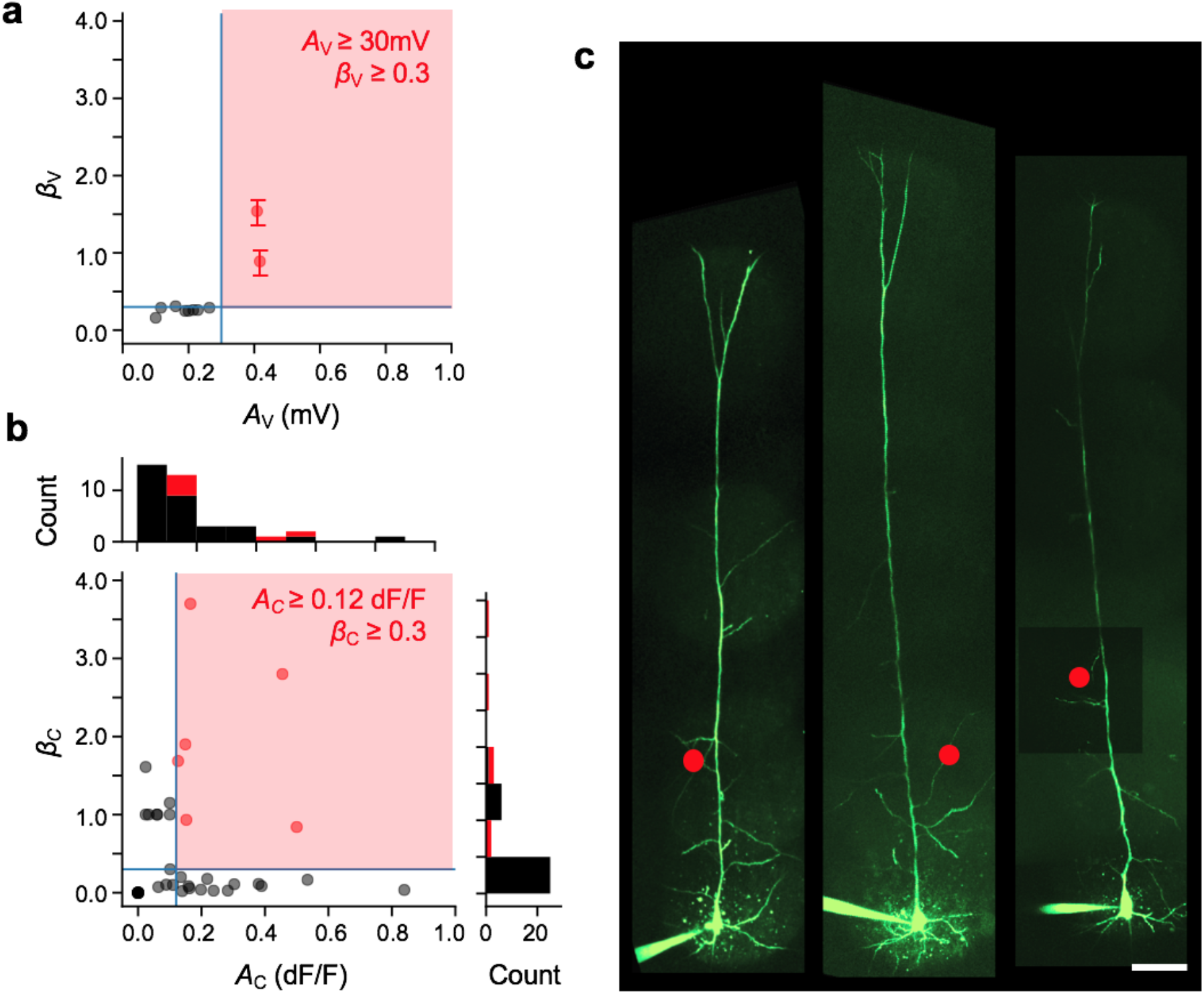
Classification of apical obliques that exhibit an oblique branch spike. (**a**) Plot of the non-linear parameter, *β*_V_, and amplitude, *A*_V_, from sigmoid fits of average membrane potential versus two-AP train frequency in oblique dendrites in the L5PN model. The value of *β*_V_ is averaged over multiple segments while the error bar shows the confidence interval (±s.d/√*N*) for *N* segments along an oblique branch. The red shaded region indicates values of *A*_V_ ≥ 30 mV and *β*_V_ ≥ 0.3. (**b**) Plot of *β*_C_ and *A*_C_ for sigmoid fits to experimentally recorded calcium transients versus two-AP train frequency in oblique dendrites (see **Figure 3e**) with the threshold region *A*_C_ ≥ 0.12 dF/F and *β*_C_ ≥ 0.3 shaded red. (**c**) Representative 2P images of L5PNs filled with Alexa-488 with the location of oblique dendrites that exhibited a branch spike indicated (red). The scale bar in Panel **c** is 50 μm.

As indicated in the model, dendritic spikes in the apical tuft as well as in oblique branches both lead to a step increase in the ADP at the soma at the critical frequency. We therefore investigated whether oblique branch spikes in L5PNs *in vitro* lead to a step increase in the ADP at the soma during two-AP trains, as occurs during apical tuft calcium spikes (see Larkum et al. 1999). To investigate this, we recorded the ADP at the soma in a large population of L5PNs (*N*=100 L5PNs) during two- and four-AP trains at different frequencies (**Figure 5a**).

**Figure 5.**
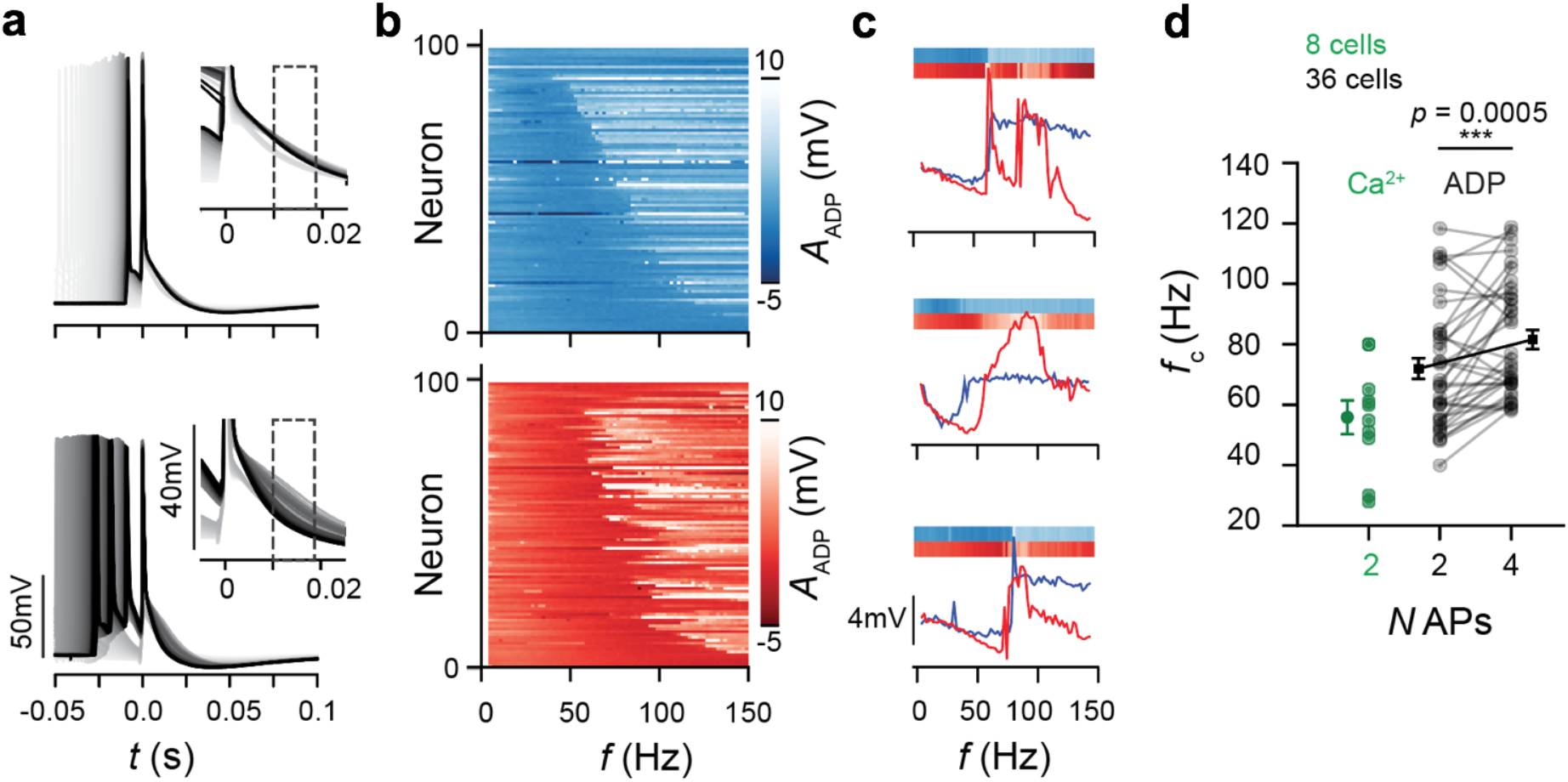
Analysis of the critical frequency from the somatic ADP during AP trains. (**a**) After depolarizing potential (ADP) in L5PNs during two- and four-AP trains at frequencies from 4 < *f* <150 Hz. Traces are aligned to the last AP (*t* = 0) of the train, with dotted box showing when the ADP was measured (inset). (**b**) Heat maps of the step increase in the average ADP amplitude for two-AP (top, blue) and four-AP trains (bottom, red) at different frequencies (*N* = 100 L5PNs). Data sorted according to the critical frequency (*f*_c_) for two-AP trains. (**c**) Sample ADP responses from three L5PNs showing variable *f*_c_’s for two (blue) and four (red) AP trains at different frequencies. (**d**) Summary plot of *f*_c_s from calcium imaging at oblique dendrites (green) and the ADP at the soma for two- and four-AP trains.

In general, there was a lot of variability in the critical frequency for generation of a step change in the ADP during both two- and four-AP trains (**Figure 5b**). Some L5PNs showed a lower critical frequency for two-AP trains than four-AP trains (**Figure 5c**, middle), while in others the critical frequency was the same for two- and four-AP trains (**Figure 5c**, bottom). To identify which L5PNs exhibited a non-linear step increase in the ADP during two- and four-AP trains, we applied the criteria for the slope (*β*_A_ ≥ 0.3), while we used an amplitude threshold *A*_A_ ≥ 1 mV. A significant fraction (27%) of all L5PNs investigated did not exhibit a detectable step-increase in the ADP (27 out of 100 L5PNs). Approximately a third (35%; 35 out of 100) showed a step increase in the ADP only for four-AP trains, with an average critical frequency of 98±4 Hz. A third group representing 36% of L5PNs (36 out of 100) displayed a step increase in the ADP for both two-AP trains at 72±4 Hz and four-AP trains at 82±3 Hz. Finally, a very small percentage of recorded L5PNs (2%; 2 out of 100) showed a step increase only during two-AP trains at 75±10 Hz. **Figure 5d** shows the summary of critical frequencies from L5PNs that showed detectable increases in the ADP during both two- and four-AP trains. The critical frequencies for two- and four-AP trains were significantly different (*p*=0.0005: paired student’s *t*-test, *N*=36 L5PNs). These results are consistent with our model and show that the step increase in the ADP occurs at two distinct critical frequencies during two- and four-AP trains (**Figure 1c**). On average, the critical frequency obtained from the ADP measurements (72±4 Hz) is comparable to the one observed during calcium imaging (58±5 Hz), consistent with the idea that the step increase in the ADP during 2-AP trains is a result of the generation of an oblique branch spike. In contrast, the critical frequency during four-AP trains (82±3 Hz) is comparable to that reported by (Larkum et al., 1999) and presumably represents generation of a dendritic calcium spike in the nexus of the apical tuft. Finally, we verified that the step increase in the ADP during 2-AP trains were not due to generation of NMDA spikes, as bath applications of the NMDA antagonist (25μM DL-APV) had no effect on non-linear increases in the ADP during two-AP trains (**Supplementary Figure 6**). These data are consistent with the idea that the oblique branch spikes we report here are intrinsically generated via the depolarization associated with low frequency AP trains.

## Discussion

Dendritic spikes enable neurons to perform complex dendritic computations on their inputs. The ionic mechanisms and conditions for generation of dendritic spikes have been well documented (Sjostrom et al., 2008, London and Hausser, 2005, Major et al., 2013, Ramaswamy and Markram, 2015, Stuart and Spruston, 2015). In this study, we investigated the conditions for generation of oblique branch spikes in a multi-compartment model and then use two-photon calcium imaging and ADP analysis in acute brain slices to confirm their existence in real neurons. We find that a subset of oblique dendrites in L5PNs can generate dendritic spikes during trains of two APs at low frequencies.

Based on simulations in the multi-compartment model of a L5PN, we established that two-AP trains at around 40 Hz elicit dendritic spikes in 2 out of 9 oblique branches. These oblique branch spikes consisted of a fast Na^+^ component followed by a broad depolarization due to the recruitment of voltage-activated calcium channels. Such low frequency AP trains were not enough to elicit a calcium spike at the nexus of the apical tuft. Similar to these findings in the multi-compartment model, we observed oblique branch spikes in L5PNs *in vitro* during low-frequency two-AP trains. Based on two-photon calcium imaging, oblique branch spikes were evoked at an average critical frequency of around 60 Hz, whereas ADP measurements at the soma yielded a critical frequency of around 70 Hz. One complication with measurement of the ADP is that changes in the ADP may result from dendritic spikes at multiple locations, including the nexus of the apical tuft (**Figure 2d**). As such, the critical frequency obtained from ADP measurements at the soma is likely to exhibit large variance during two- and four-AP trains, as observed from the heat maps in **Figure 5b**. The difference in the critical frequencies obtained from calcium imaging and the ADP analysis could also reflect ineffective propagation of some oblique branch spikes to the soma, possibly at dendritic branch points. In general, the critical frequency determined experimentally was higher than in the model. This difference could be due to differences in activation of dendritic A-type potassium channels, which have been reported to dampen the invasion of APs into oblique branches in hippocampal CA1 neurons (Frick et al., 2003, Gasparini et al., 2007).

### Oblique branch spikes in hippocampal neurons

Dendritic spikes in oblique branches of hippocampal CA1 pyramidal neurons have been studied extensively (Kamondi et al., 1998, Losonczy and Magee, 2006, Canepari et al., 2007). Dendritic recordings from radial oblique branches of CA1 pyramidal neurons indicate they can evoke large and fast spikes, including putative calcium spikes (Kamondi et al., 1998). Losonczy and Magee (2006) demonstrated that clustered activation of synapses in oblique branches also triggers dendritic Na^+^ spikes whose temporal profile is shaped by coincident activation of NMDA receptors, voltage-activated calcium channels, and transient potassium currents. In addition, pairing EPSPs and APs leads to supra-linear Ca^2+^ signals in the oblique dendrites of CA1 PNs (Canepari et al., 2007). While some of the properties of oblique dendrites of cortical L5PNs are likely to be similar to oblique dendrites in CA1 pyramidal neurons, the oblique branch spikes triggered by low frequency trains of APs in L5PNs have not yet been observed in CA1 pyramidal neurons.

### Branch specific nature of dendritic oblique spikes

The dendritic spikes we report here only occurs in a subset of oblique dendrites. A possible explanation for this branch specificity is that the efficacy of AP backpropagation into oblique dendrites is heterogenous, due to differences in dendritic morphology and/or ion channel expression. Studies using multi-compartmental models indicate that attenuation of bAPs is dependent on multiple factors, such as the extent of dendritic branching, the number of branch points, branch-point morphology, dendritic diameter, and presence of leak currents (Vetter et al., 2001, Schaefer et al., 2003, Migliore et al., 2005, Ferrante et al., 2013). *In vitro* patch-clamp recordings and voltage imaging in L5PNs confirm that AP backpropagation is highly dependent on activation of voltage-dependent channels and morphology as well as distance from the soma (Stuart et al., 1997b, Waters et al., 2005, Major et al., 2013). Aside from branching patterns, other studies have shown that the heterogeneous distribution of A-type potassium channels, which opposes membrane excitability, impact on AP invasion into the apical tuft and apical oblique dendrites (Hoffman et al., 1997, Frick et al., 2003, Gasparini et al., 2007, Zhou et al., 2015, Short et al., 2017). Future experiments that are capable of manipulating the morphology (dendrotomy) and activation of voltage-dependent channels in specific oblique branches may shed light onto what underlies why oblique branch spikes were observed in some, but not all, oblique dendrites in L5PNs. In terms of function, oblique branch spikes may provide a mechanism to significantly raise intracellular calcium levels or modify expression of voltage-gated potassium channels in certain oblique branches, promoting branch-specific plasticity (Alkon, 1999, Losonczy et al., 2008).

### Future perspectives and conclusions

It will also be important to understand whether oblique branch spikes play a role in learning and memory. The critical frequency for generation of oblique branch spikes could be relevant to the induction of long-term potentiation (LTP) at proximal L5PN-L5PN synaptic connections. Sjostrom and Hausser (2006) reported that LTP was induced in paired recordings from L5PNs using an AP-EPSP pairing protocol. Interestingly, they used a low frequency AP train (50 Hz) applied extracellularly to induce LTP. Such AP trains may have recruited dendritic spikes in oblique dendrites leading to induction of LTP.

*In vivo* experiments could be performed to explore the relevance of oblique branch spikes to sensory processing. Several works have shown that dendritic Ca^2+^-AP at the nexus and NMDA in apical tuft branches enhance sensory processing (Lavzin et al., 2012, Xu et al., 2012, Smith et al., 2013, Palmer et al., 2014, Ranganathan et al., 2018). New technologies allowing imaging in deeper cortical layers could be used to optically record oblique branch spikes *in vivo* and thereby investigate their relevance to sensory processing and plasticity.

In conclusion, we show that dendritic spikes are evoked in select apical oblique dendrites in L5PNs during two-AP trains at frequencies of around 60 Hz; a frequency that is significantly lower than that required to elicit calcium spikes at the nexus of the apical tuft (Larkum et al., 1999) and in basal dendrites of L5PNs (Kampa and Stuart, 2006). The occurrence of dendritic spikes in a subset of oblique branches offers new insights into branch-specific computation in L5PNs and may be critical for sensory processing in the input layers of the cortex.

## Methods

### Multi-compartment model

We used the Shai et al. (2015) L5PN model (MODELDB 180373-cell1.asc) obtained from SenseLab Database (http://senselab.med.yale.edu/ModelDB/) to numerically investigate the active properties of oblique dendrites. We implemented a script in Neuron-Python environment (Hines et al., 2009) that simulated the critical frequency protocol and obtained the temporal profiles of the membrane potential at different dendritic segments during AP trains of different frequencies. We also performed numerical blocking experiments where the densities of voltage-gated sodium and calcium channels in oblique dendrites were set to zero. Apart from the numerical blocking experiments, the densities of all other ion-channels and their distributions along the dendritic tree were unchanged from that reported by Shai et al. (2015) (see **Supplementary Figure 1**). We labelled oblique branches with a number, *O*#*n*, which corresponded to the position of the branch from the soma (e.g., *O*#1 is the first oblique branch from the soma).

### Electrophysiology

Wistar male rats (P26–P34 of age) were sedated by inhalation of 2-4% isoflurane with 3L/min flow rate of oxygen and decapitated according to protocols approved by the Animal Experimentation Ethics Committee of the Australian National University. Somatosensory cortical brain slices (300-μm-thick) were prepared using a vibratome and perfused with oxygenated (95% O_2_/5% CO_2_) extracellular solution containing (in mM): 125 NaCl, 25 NaHCO_3_, 25 glucose, 3 KCl, 1.25 NaH_2_PO_4_, 2 CaCl_2_, 1 MgCl_2_, pH 7.4. For patch-clamp recording and 2P imaging, tissue slices were transferred to a custom-built 2P microscope. Whole-cell recordings were made from the soma of L5PNs with 4-6 MΩ pipettes filled with an intracellular solution containing (in mM): 115 K-gluconate, 20 KCl, 10 HEPES, 10 phosphocreatine, 4.0 ATP-Mg, and 0.3 GTP (Castanares et al., 2019). During recording slices were kept at 33±1 °C with the temperature of the extracellular solution regulated by a proportional integral derivative controller (Inkbird ITC-100).

Calcium imaging was performed using the calcium indicator Cal-520 (potassium salt; AAT-Bioquest). Recording pipettes were front-filled with a dye-free intracellular solution and then back-filled with the same solution to which we added 0.3mM of Cal-520 and 0.13mM Alexa-Fluor 488 (Sigma Aldrich). Alexa-Fluor 488 was included in the internal solution to aid visualization of dendrites for two-photon imaging. Recordings were made from the soma of L5PNs using the whole-cell configuration and held at the resting potential, *V*_m_ = −65 mV. After obtaining the whole-cell configuration the recording was left undisturbed for 45-60 min to allow sufficient loading of the dye solution into the dendrites via passive diffusion. All recordings had pipette series resistances of <30 MΩ. In cases where the pipette series resistance increased beyond 30 MΩ, the cell was re-patched with a fresh pipette.

### Two-photon holographic calcium imaging

We used a custom-built two-photon microscope as previously reported (Go et al., 2013, Go et al., 2016) and modified it to include a widefield imaging camera for holographic calcium imaging (Castanares et al., 2016, Castanares et al., 2019) (**Supplementary Figure 2**). The microscope uses a near-infrared femtosecond laser (Chameleon, Coherent Scientific) and galvanometer scanning mirrors to render a 3D image of the patched neuron (**Supplementary Figure 7a**). After mapping the 3D morphology of the entire neuron, a phase-only hologram was calculated to target dendrites of interest. A fraction of the incident laser (via a polarizing beam splitter, PBS) was expanded and directed towards a spatial light modulator (SLM, Hamamatsu X10468-02) where the phase-only hologram is encoded. The hologram-encoded laser from the SLM was then directed towards the brain tissue. The hologram transforms the incident laser into multiple foci allowing two-photon excitation simultaneously at multiple dendritic locations (**Supplementary Figure 7b**). The fluorescence from multiple foci were simultaneously acquired using an electron-multiplying charged coupled camera (EMCCD, Andor Ixon 897, Oxford Instruments) at 100 frames per second. Since the imaging plane is fixed, we selected oblique dendrites that ran parallel to the imaging plane based on the 3D image.

### Critical frequency protocol

Two- and four-AP trains at different frequencies were evoked by current pulses (3-4 nA, 2 ms) injected into the soma (Larkum et al., 1999). Sweeping the frequency of AP trains over a range (15 < *f* < 200 Hz) allowed the critical frequency, *f*_c_, for generation of dendritic spikes to be determined. The *f*_c_ is the minimum frequency of an AP train that evokes a dendritic spike. We applied the protocol in both multi-compartmental model and *in vitro* experiments. In the model, we identified the critical frequency by plotting the average membrane potential over a constant time interval (120 ms) at different frequencies. For *in vitro* experiments, we measured calcium transients at specific dendritic locations during AP trains of different frequency. In addition to these dendritic measurements, we also obtained the average membrane potential at the soma during the after-depolarizing potential (ADP) that following an AP train. Plots of the average membrane potential, change in fluorescence and somatic average ADP versus AP train frequency were fitted with a sigmoid function, *S*(*f*) = *A*/(1 + exp [*β*(*f* – *f*_c_)]), where the fitting parameters *A* and *β* refer to the amplitude and slope, respectively.

### Average membrane potential and average ADP

The average membrane potential was derived from the integral of the membrane potential (*V*_m_) over a constant time interval (*t*_1_ ≤ *t* ≤ *t*_2_) and follows the equation: Average 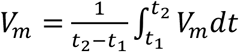. To obtain the average membrane potential during a two-AP train (*in silico*), the membrane potential was integrated from the start of the simulation run, *t*_1_=0 to *t*_2_=120 ms. The time interval ensures that the two-AP train with frequencies *f* ≥ 15 Hz is contained within the time window.

The average ADP measured *in silico* and experimentally *in vitro* was obtained by first aligning the somatic membrane potential with the last AP of the train, which was set to be *t* = 0. Next, we calculated the average ADP for a given time window. For the L5PN model, we used the time-windows: *t*_1_=10 ms to *t*_2_=15 ms (for two-AP trains) and *t*_1_=25 ms to *t*_2_=30 ms (for four-AP trains). For *in vitro* L5PN recordings, we used the time-window of *t*_1_=10 ms to *t*_2_=18 ms (for two- and four-AP trains). Prior to measuring the average ADP amplitude, we subtracted average ADP for the lowest frequency train (*f*=15 Hz) to remove the offset in ADP-frequency scatter plots.

### Calcium responses at the oblique branches

We investigated AP-associated calcium responses in different oblique branches using the critical frequency protocol with two-AP trains. AP train frequency was randomized. Calcium transients were derived from the change in fluorescence intensity divided by the resting fluorescence intensity (dF/F). We acquired calcium transients during single APs or two-AP trains, termed *C*_1_ and *C*_2_, respectively, at one-minute intervals to allow for the intracellular calcium concentration to recover back to baseline (see **Supplementary Figure 8**). We subtracted the calcium transient evoked by a AP from that evoked by the two-AP train to yield Δ*C*_21_ = *C*_2_-*C*_1_. To determine the critical frequency the amplitude of Δ*C*_21_ was plotted versus the two-AP train frequency and the data fitted with a sigmoid.

### Statistics

For **Figures 3 and 5**, the average values of the critical frequencies represent the mean ± the S.E.M. For **Supplementary Figure 5c**, paired student t-tests were used to assess statistically significant differences between average values of *A*_c_ below and above the critical frequency for a given *p*-value (*p*<0.10, * *p*<0.05, ** *p*<0.01, ***).

## Acknowledgements

This work was primarily funded by the National Health and Medical Research Council (PG1105944) and supported by the Australian Research Council (ARC) Discovery Project (DP140101555) and the ARC Centre of Excellence for Integrative Brain Function (CIBF).

## Supplementary information

**Supplementary Figure 1.**
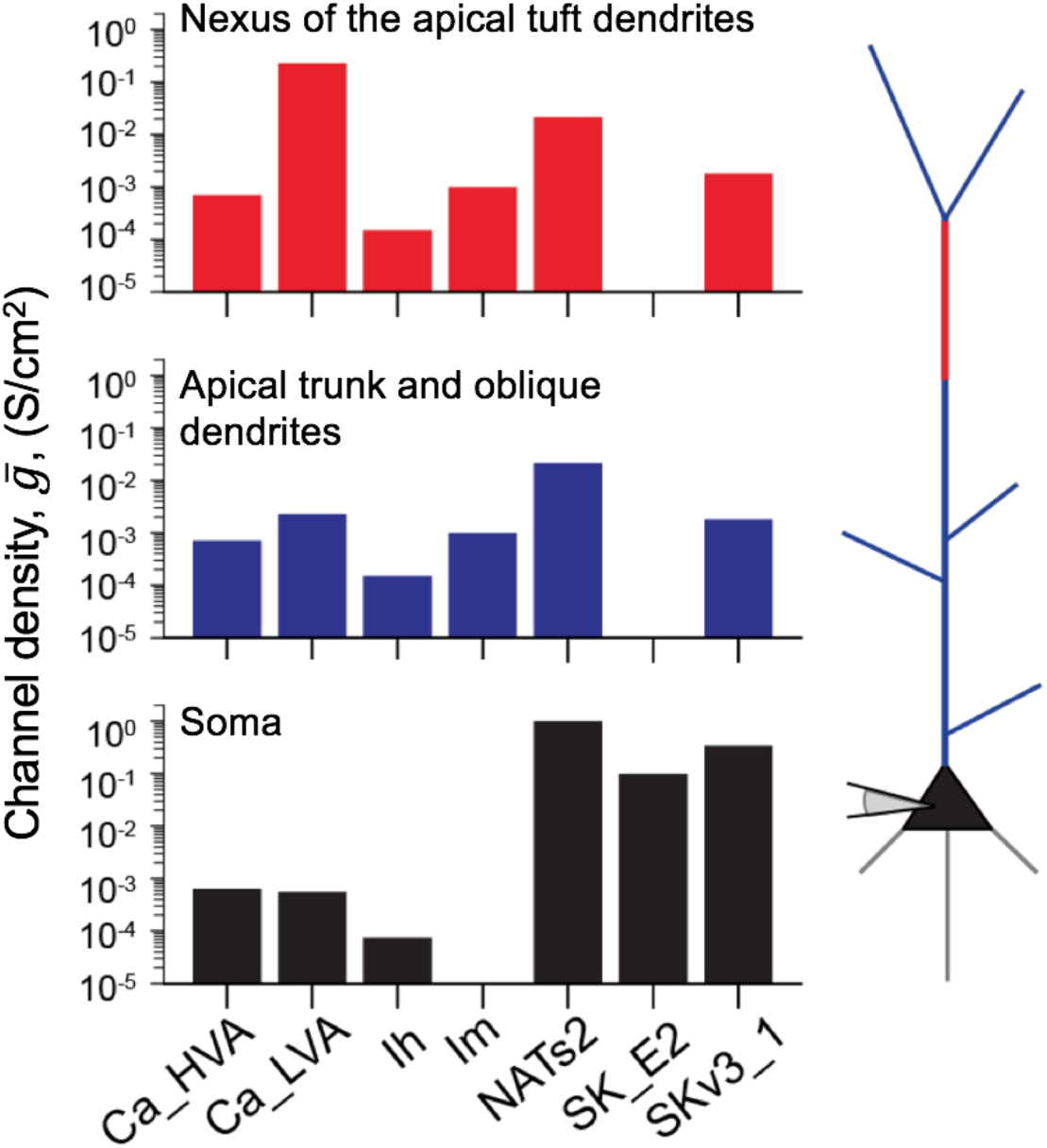
The distribution of voltage-gated and calcium activated ion channels in the Shai. et al (2015) L5PN model. The channel densities of high voltage activated 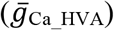, low-voltage activated 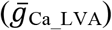 calcium channels and sodium channels 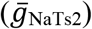 were constant along the apical trunk and oblique branches. There was a high magnitude of 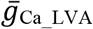 along the nexus of the apical trunk (trunk segments located 685-885 μm from the soma) to model the Ca^2+^-AP spike.

**Supplementary Figure 2.**
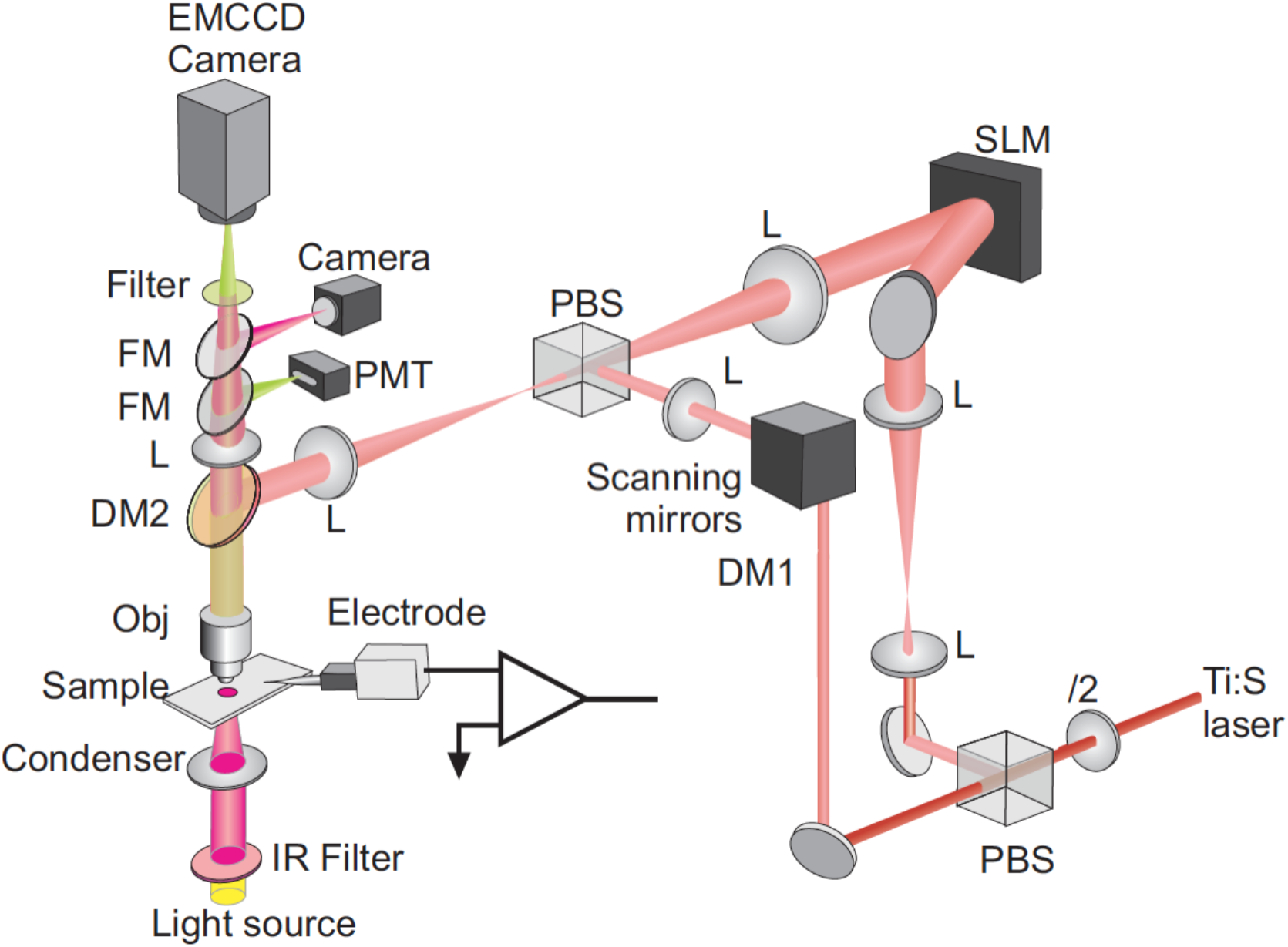
Schematic diagram of our custom-built 2P laser scanning and holographic microscope. The NIR Ti:S femtosecond laser (MIRA 900, Coherent Inc.) pumped by 12 W optically pumped semiconductor laser (Verdi G, Coherent Inc.) was directed by a polarizing beam splitter (PBS) to the laser scanning arm or the holographic arm by adjusting the polarization angle of the beam relative to the PBS. In laser scanning mode, the beam is deflected using galvanometer mirrors (Thorlabs) and the fluorescence is collected using a PMT. In holographic mode, the beam encoded with a phase map via a spatial light modulator (LCOS-based phase only SLM, Hamamatsu) and relayed via a 4f-lens configuration (FTL and TL) and a dichroic mirror (λ=805 nm short-pass, DMSP805, Thorlabs) to the back aperture of the objective lens (Obj, 40x, NA=1.0) of an upright differential interference contrast (DIC) microscope (BX50WI, Olympus). Along with the microscope is a patch-clamp system consisting of micromanipulators (Sutter Instruments) and a current amplifier (Multi-clamp 700B, Molecular Devices). The beam is shaped into user-positioned multi-foci at the sample. The incident beam is blocked with an optical filter (Filter, 810nm short-pass, Thorlabs). The fluorescence of each site is simultaneously recorded by an electron-multiplying charged coupled device (EMCCD) camera (iXon Ultra 897, Andor, Oxford Instruments).

**Supplementary Figure 3.**
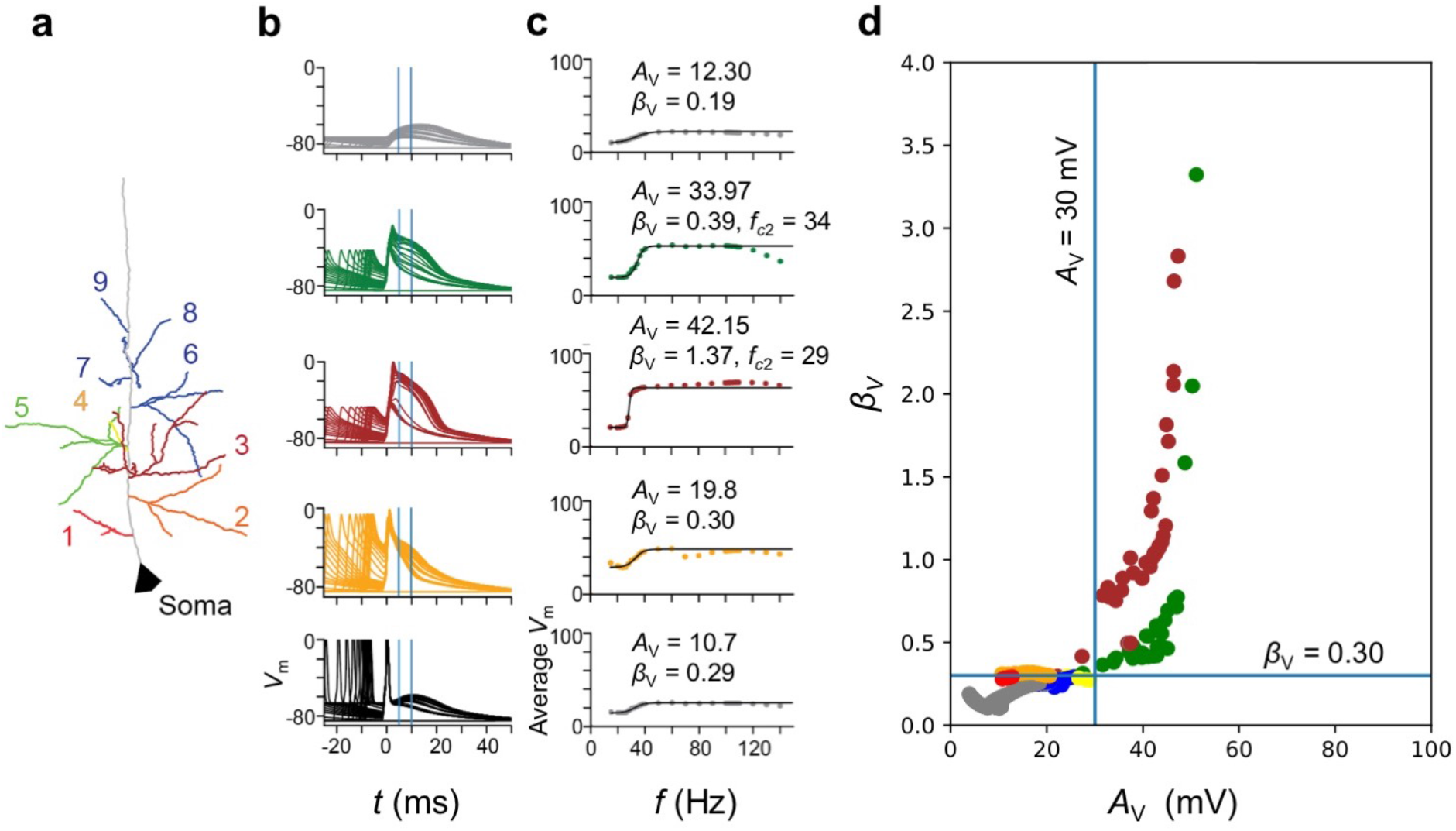
Analysis of segments that exhibited an oblique branch spike. (**a**) The apical oblique branches labelled according to the branch number, *O*#*N*. The color assignments are: soma (black), O#1 (red), O#2 (orange), O#3 (brown), O#4 (yellow), O#5 (green), O#6 to O#9 (blue), and main apical trunk (grey). (**b**) The recorded membrane potential of the segments (i.e., soma, O#2, O#3, O#5, and apical trunk) in with the firing of two-AP train with frequency ranging from 15 < *f* < 140 Hz. The dendritic recordings are aligned to the peak of the 2^nd^ AP at the soma (*t*=0). (**c**) The plot of the average *V*_m_, within the time window (*t*_0_ = 5ms, *τ* = 5ms, indicated by blue lines in **b**) versus the frequency of the two-AP train. The data was fitted with a sigmoid function and the fit parameters *A*_V_ and *β*_V_ are shown in the inset. Also, for data that exhibited (*A*_V_ ≥ 20mV and *β*_V_ ≥ 0.3), the *f*_c2_ is also indicated. (**c**) The plot of *β*_V_ versus *A*_V_ with the points color-coded according to dendritic segment they belong (i.e., apical oblique dendrites or main apical trunk).

**Supplementary Figure 4.**
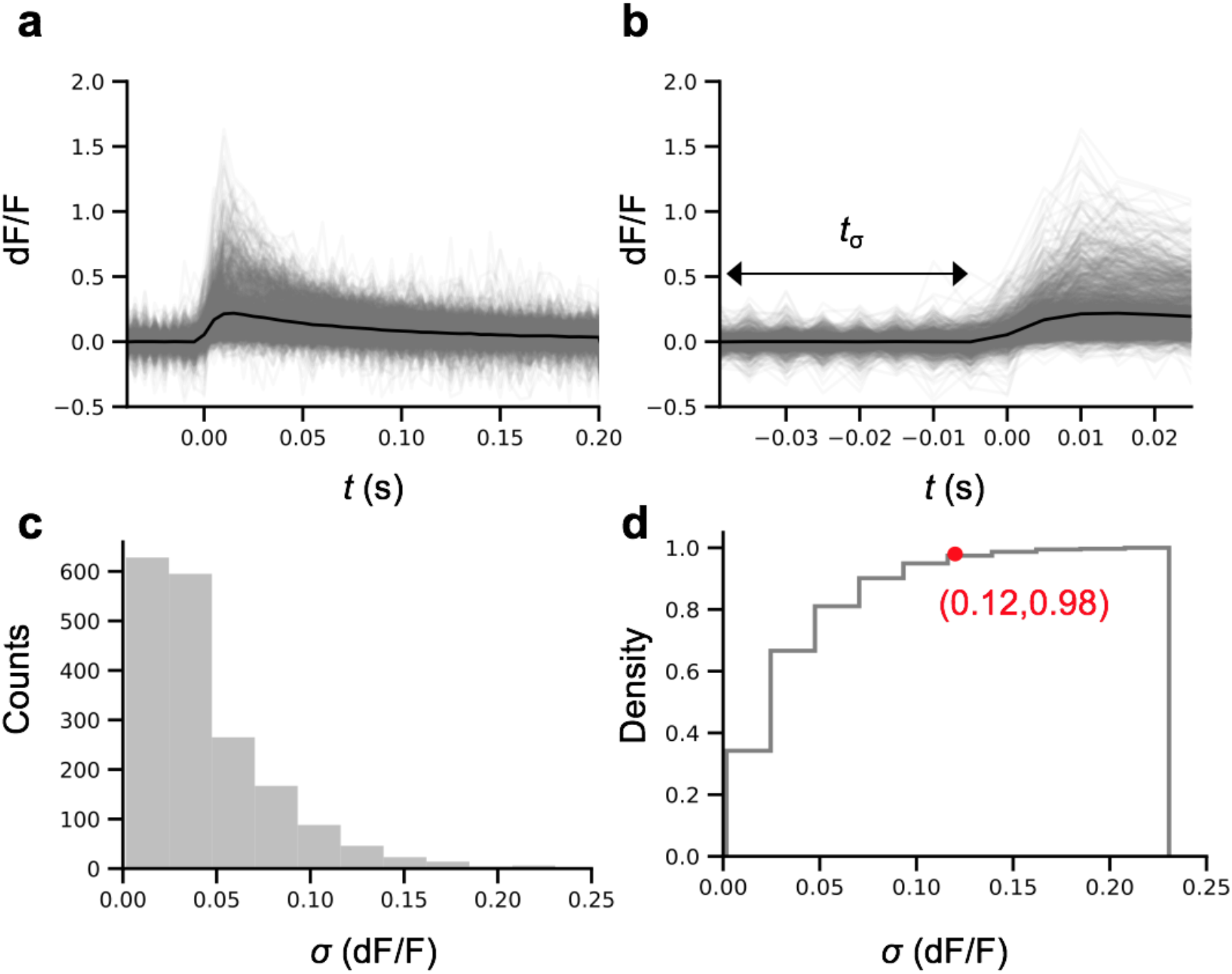
Estimation of signal to noise of the recorded calcium transients. (**a**) The relative changes in the fluorescence recorded from oblique dendrites (*N* = 1794 total traces taken from each AP-train frequency from 11 L5PNs, black trace is the average of all the traces) associated with single and two-AP firing. (**b**) An expanded view of the initial part of the calcium transient shown in (**a**), with the time window, *t*_σ_, over which the baseline noise was measured. (**c**) A histogram of the standard deviation of the noise from the recorded calcium transients. (**d**) The cumulative histogram of the standard deviation of the noise of the calcium transients. The red dot indicates where 0.12 dF/F covered 98% of the distribution.

**Supplementary Figure 5.**
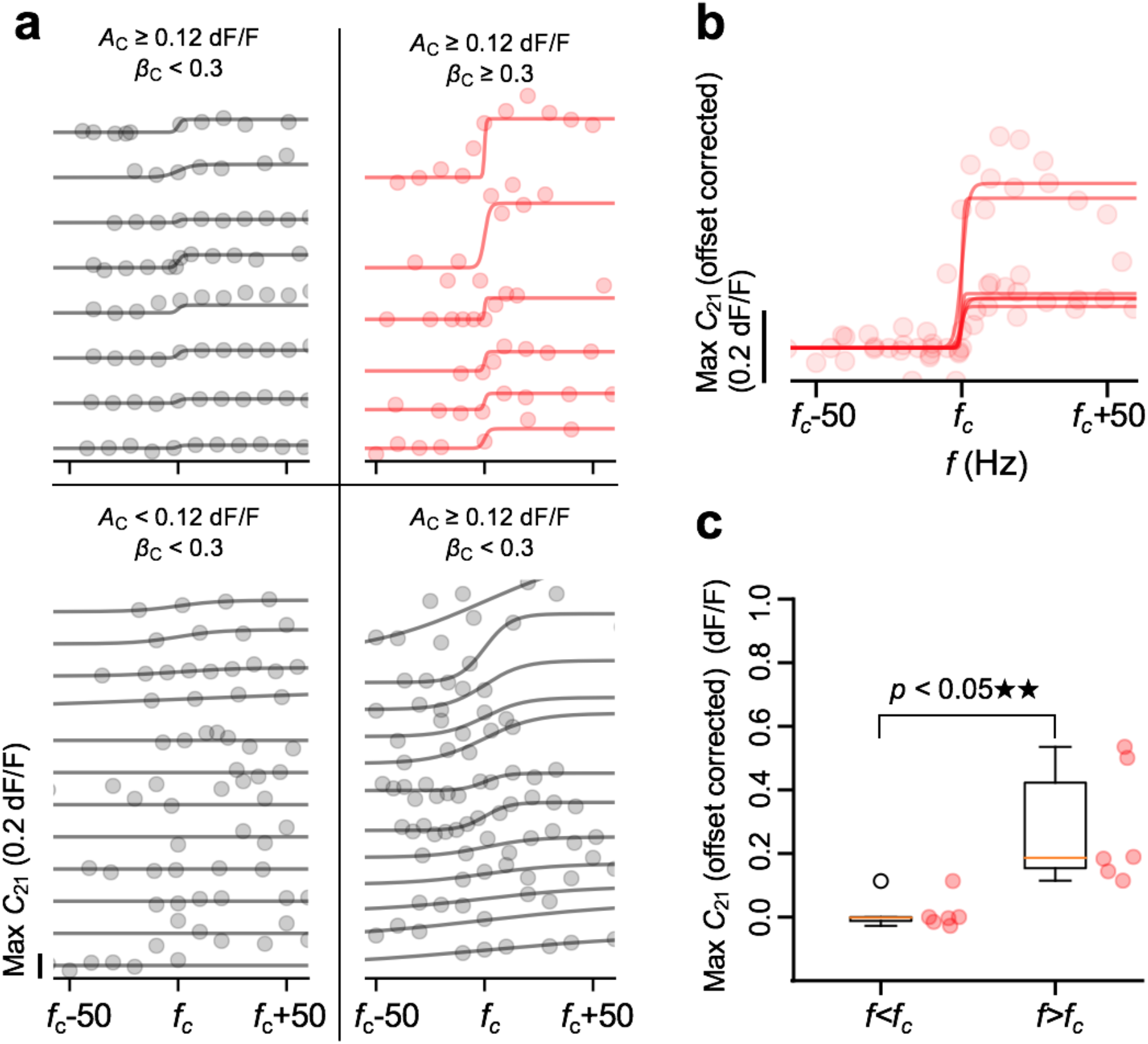
Calcium response versus AP train frequency for different L5PNs oblique dendrites. (**a**) Plots of the amplitude of calcium responses versus AP train frequency (*f*_c_) for all recorded L5PNs oblique dendrites (*N*=38) separated into 4 quadrants based on *β*_c_ values greater than or less than 0.3 (left, top & bottom, respectively) and *A*_c_ values greater than or less than 0.12 dF/F (right & left, respectively). (**b**) Overlaid offset-corrected maximum calcium response amplitude versus frequency plots for oblique dendrites where *A*_c_ ≥ 0.12 and *β* ≥ 0.3, indicating generation of an oblique branch spike. (**c**) A paired *t*-test of Max. *C*_21_ (offset corrected) between frequencies below (*f*<*f*_c_) and above (*f*>*f*_c_) the critical frequency (*t* = −3.79, *p* = 0.0128, *N* = 6 L5PNs, df = 5).

**Supplementary Figure 6.**
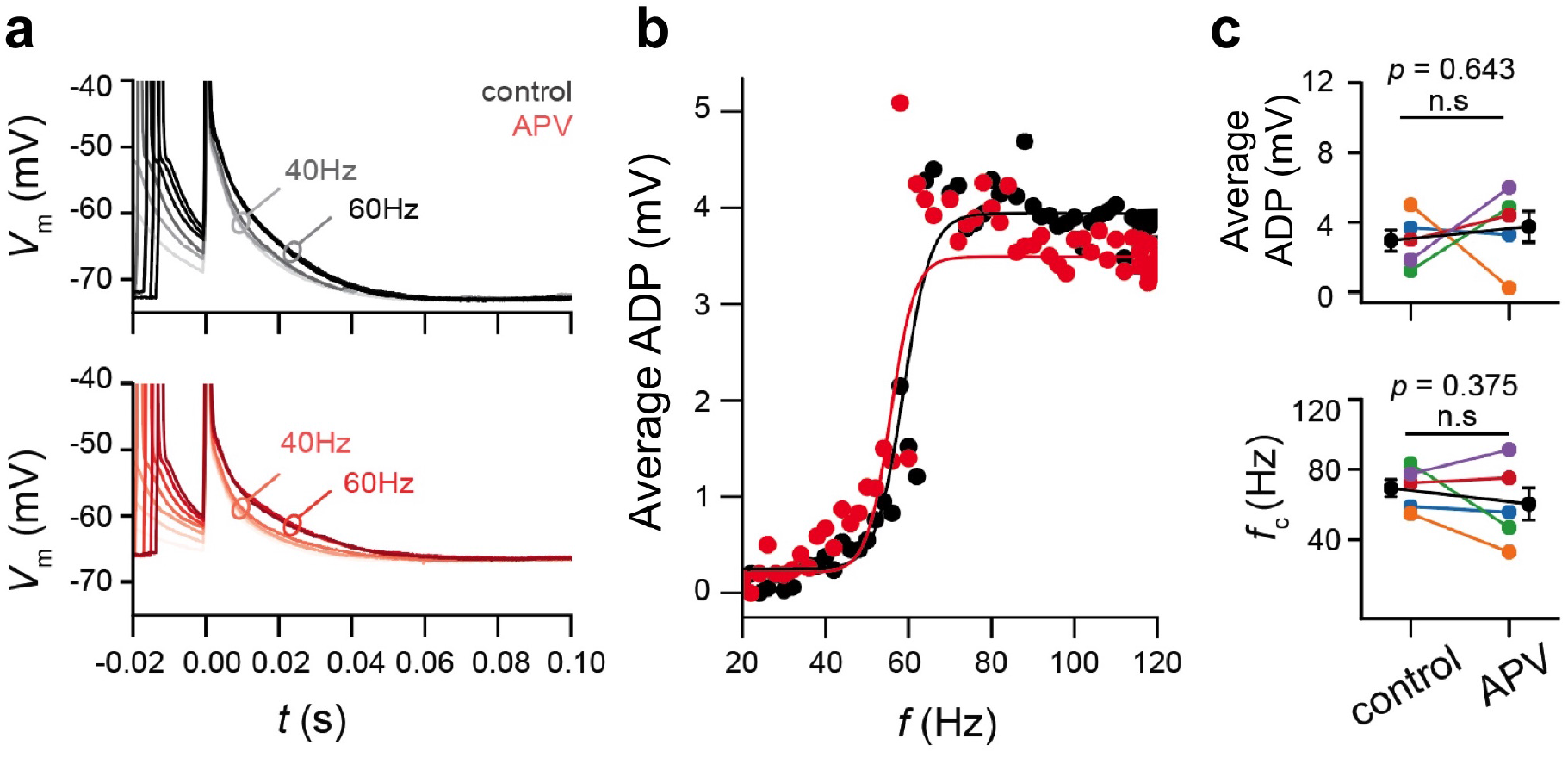
(**a**) Somatic *V*_ADP_ recording in a L5PN bathed in ACSF solution (control) and after 15 min bath application of 25 μM APV (an NMDA antagonist). (**b**) The plot of the average *V*_ADP_ with *f* of the recordings in (**a**). (**c**) A summary plot of the sigmoid fit parameters of the ΔADP with *f* from sample L5PNs (*N* = 5) under control and APV with their *p* – values for average ADP (student’s paired t-test two-tailed, t = 2.77, *p* = 0.643, *N* = 5, df = 4) and for critical frequency (student’s paired t-test two-tailed, t = 2.77, *p* = 0.375, *N* = 5, df = 4).

**Supplementary Figure 7.**
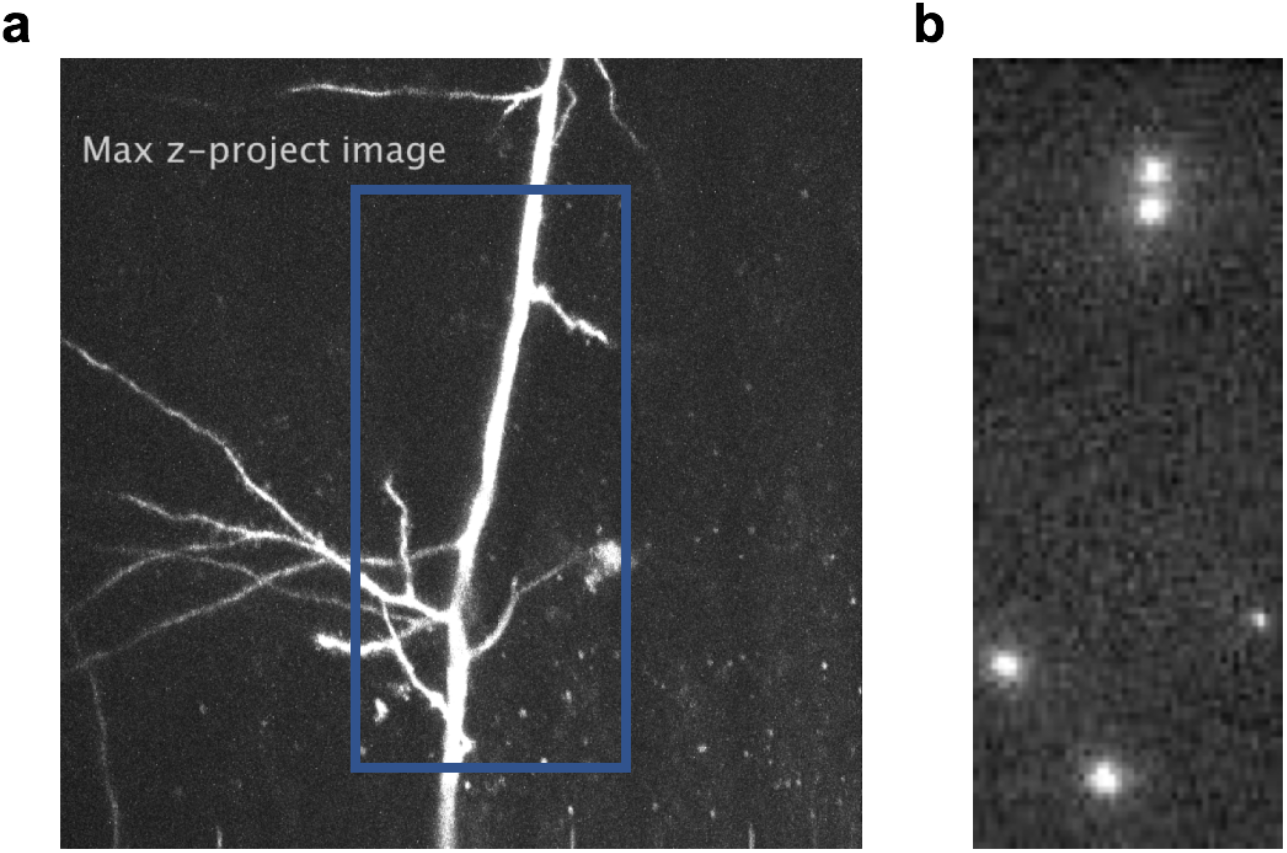
A sample z-stack 2P images of an oblique branch and the fluorescence recording of from the holographic sites. (**a**) A z-stack images of an oblique dendrite branching in 3D space. The 2D z-projection image generates a simplified view of the branching pattern. (**b**) An image sequence of the calcium fluorescence transients from 5 holographically projected foci at the oblique branches and the apical trunk of an L5PN.

**Supplementary Figure 8.**
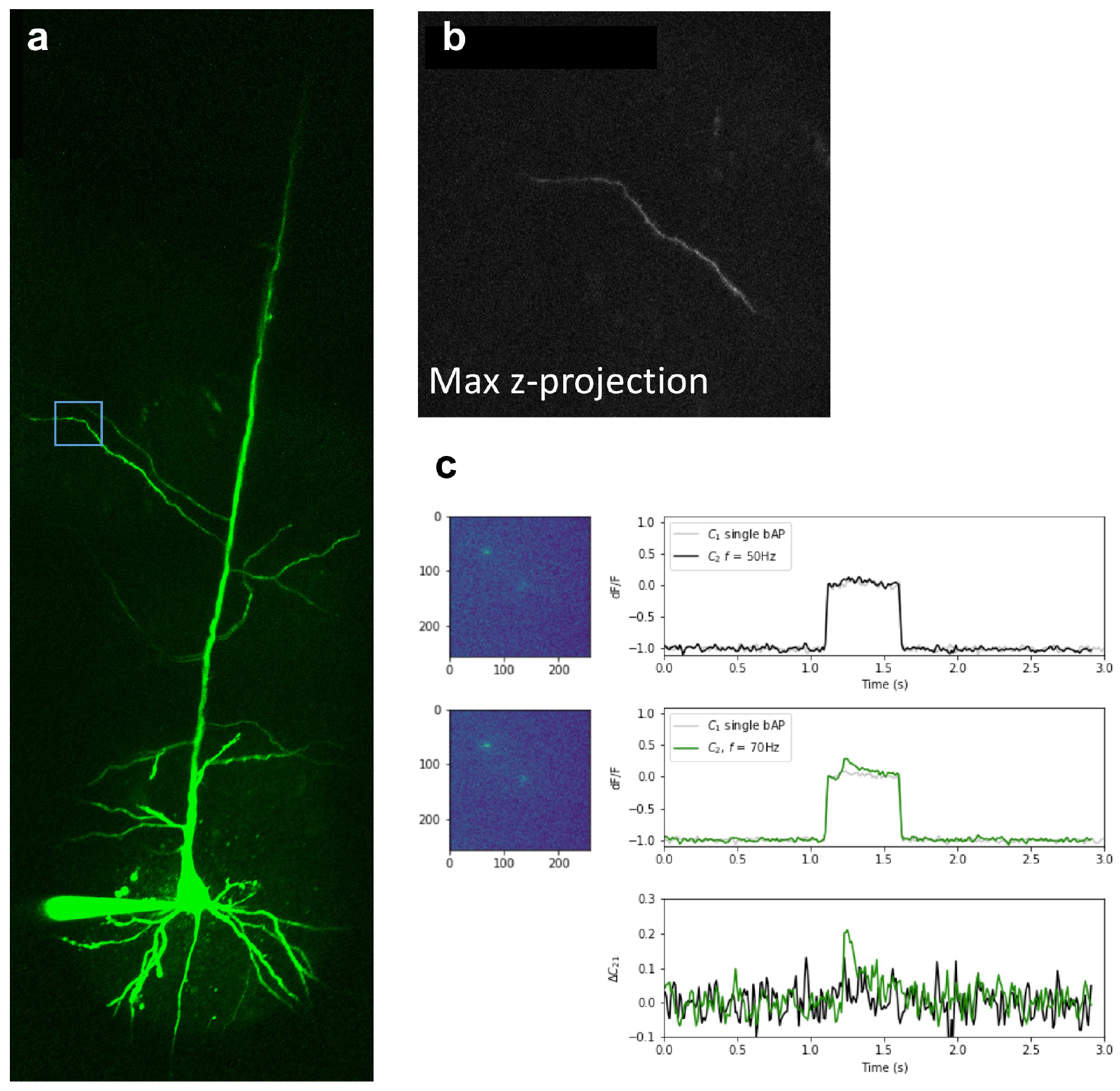
A sample z-stack 2P images of an oblique branch and the fluorescence recording of from the holographic sites. (**a**) A stitched image reconstructing the dendritic tree of an L5PN loaded with Alexa-488 fluorescence indicator and Cal-520 calcium indicator with the 2P laserscanning field-of-view (white box) and the EMCCD’s active recording pixels (256×256 pixels, blue box). (**b**) A z-stack images of an oblique dendrite branching in 3D space. The 2D z-projection image generates a simplified view of the branching pattern. (**c**) The raw frame capture of the fluorescence from 3 holographically projected foci at an oblique branch (left) and the average intensity of the response with time (right). The images were captured at 100 frames/second. For comparison, the dF/F responses for two-AP trains with *f* = 50 Hz (top) and *f* = 70 Hz (middle) are shown. (bottom) In addition, the relative increase in the calcium transients, Δ*C*_21_, are also shown for *f* = 50 Hz (black) and *f* = 70 Hz (green), respectively.

## References

Alkon, D. L. 1999. Ionic conductance determinants of synaptic memory nets and their implications for Alzheimer’s disease. J Neurosci Res, 58, 24–32.

Amitai, Y., Friedman, A., Connors, B. W. & Gutnick, M. J. 1993. Regenerative activity in apical dendrites of pyramidal cells in neocortex. Cereb Cortex, 3, 26–38.

Antic, S. D. 2003. Action potentials in basal and oblique dendrites of rat neocortical pyramidal neurons. Journal of Physiology-London, 550, 35–50.

Beaulieu-Laroche, L., Toloza, E. H. S., Brown, N. J. & Harnett, M. T. 2019. Widespread and Highly Correlated Somato-dendritic Activity in Cortical Layer 5 Neurons. Neuron, 103, 235–241 e4.

Beaulieu-Laroche, L., Toloza, E. H. S., Van Der Goes, M. S., Lafourcade, M., Barnagian, D., Williams, Z. M., Eskandar, E. N., Frosch, M. P., Cash, S. S. & Harnett, M. T. 2018. Enhanced Dendritic Compartmentalization in Human Cortical Neurons. Cell, 175, 643–651 e14.

Branco, T., Clark, B. A. & Hausser, M. 2010. Dendritic discrimination of temporal input sequences in cortical neurons. Science, 329, 1671–5.

Canepari, M., Djurisic, M. & Zecevic, D. 2007. Dendritic signals from rat hippocampal CA1 pyramidal neurons during coincident pre- and post-synaptic activity: a combined voltage- and calcium-imaging study. J Physiol, 580, 463–84.

Castanares, M. L., Gautam, V., Drury, J., Bachor, H. & Daria, V. R. 2016. Efficient multi-site two-photon functional imaging of neuronal circuits. Biomedical Optics Express, 7, 5325–5334.

Castanares, M. L., Stuart, G. J. & Daria, V. R. 2019. Holographic Functional Calcium Imaging of Neuronal Circuit Activity. In: Kao, F.-J., Keiser, G. & Gogoi, A. (eds.) Advanced Optical Methods for Brain Imaging. Singapore: Springer.

Constantinople, C. M. & Bruno, R. M. 2013. Deep cortical layers are activated directly by thalamus. Science, 340, 1591–4.

Ferrante, M., Migliore, M. & Ascoli, G. A. 2013. Functional Impact of Dendritic Branch-Point Morphology. Journal of Neuroscience, 33, 2156–2165.

Fletcher, L. N. & Williams, S. R. 2019. Neocortical Topology Governs the Dendritic Integrative Capacity of Layer 5 Pyramidal Neurons. Neuron, 101, 76–90 e4.

Frick, A., Magee, J., Koester, H. J., Migliore, M. & Johnston, D. 2003. Normalization of Ca2+ signals by small oblique dendrites of CA1 pyramidal neurons. Journal of Neuroscience, 23, 3243–3250.

Gasparini, S., Losonczy, A., Chen, X., Johnston, D. & Magee, J. C. 2007. Associative pairing enhances action potential back-propagation in radial oblique branches of CA1 pyramidal neurons. J Physiol, 580, 787–800.

Gidon, A., Zolnik, T. A., Fidzinski, P., Bolduan, F., Papoutsi, A., Poirazi, P., Holtkamp, M., Vida, I. & Larkum, M. E. 2020. Dendritic action potentials and computation in human layer 2/3 cortical neurons. Science, 367, 83–87.

Go, M. A., Choy, J. M. C., Colibaba, A. S., Redman, S., Bachor, H. A., Stricker, C. & Daria, V. R. 2016. Targeted pruning of a neuron’s dendritic tree via femtosecond laser dendrotomy. Scientific Reports, 6.

Go, M. A., To, M. S., Stricker, C., Redman, S., Bachor, H. A., Stuart, G. J. & Daria, V. R. 2013. Four-dimensional multi-site photolysis of caged neurotransmitters. Frontiers of Cellular Neuroscience, 7, 231.

Harnett, M. T., Magee, J. C. & Williams, S. R. 2015. Distribution and function of HCN channels in the apical dendritic tuft of neocortical pyramidal neurons. J Neurosci, 35, 1024–37.

Hines, M. L., Davison, A. P. & Muller, E. 2009. NEURON and Python. Front Neuroinform, 3, 1.

Hoffman, D. A., Magee, J. C., Colbert, C. M. & Johnston, D. 1997. K+ channel regulation of signal propagation in dendrites of hippocampal pyramidal neurons. Nature, 387, 869–75.

Kamondi, A., Acsady, L. & Buzsaki, G. 1998. Dendritic spikes are enhanced by cooperative network activity in the intact hippocampus. J Neurosci, 18, 3919–28.

Kampa, B. M. & Stuart, G. J. 2006. Calcium spikes in basal dendrites of layer 5 pyramidal neurons during action potential bursts. Journal of Neuroscience, 26, 7424–7432.

Kim, H. G. & Connors, B. W. 1993. Apical dendrites of the neocortex: correlation between sodium- and calcium-dependent spiking and pyramidal cell morphology. J Neurosci, 13, 5301–11.

Larkum, M. E., Kaiser, K. M. M. & Sakmann, B. 1999. Calcium electrogenesis in distal apical dendrites of layer 5 pyramidal cells at a critical frequency of back-propagating action potentials. Proceedings of the National Academy of Sciences of the United States of America, 96, 14600–14604.

Larkum, M. E., Nevian, T., Sandler, M., Polsky, A. & Schiller, J. 2009. Synaptic integration in tuft dendrites of layer 5 pyramidal neurons: a new unifying principle. Science, 325, 756–60.

Larkum, M. E., Waters, J., Sakmann, B. & Helmchen, F. 2007. Dendritic spikes in apical dendrites of neocortical layer 2/3 pyramidal neurons. JNeurosci, 27, 8999–9008.

Lavzin, M., Rapoport, S., Polsky, A., Garion, L. & Schiller, J. 2012. Nonlinear dendritic processing determines angular tuning of barrel cortex neurons in vivo. Nature, 490, 397–401.

London, M. & Hausser, M. 2005. Dendritic computation. Annu Rev Neurosci, 28, 503–32.

Losonczy, A. & Magee, J. C. 2006. Integrative properties of radial oblique dendrites in hippocampal CA1 pyramidal neurons. Neuron, 50, 291–307.

Losonczy, A., Makara, J. K. & Magee, J. C. 2008. Compartmentalized dendritic plasticity and input feature storage in neurons. Nature, 452, 436–41.

Major, G., Larkum, M. E. & Schiller, J. 2013. Active properties of neocortical pyramidal neuron dendrites. Annu Rev Neurosci, 36, 1–24.

Markram, H., Lubke, J., Frotscher, M., Roth, A. & Sakmann, B. 1997. Physiology and anatomy of synaptic connections between thick tufted pyramidal neurones in the developing rat neocortex. J Physiol, 500 (Pt 2), 409–40.

Migliore, M., Ferrante, M. & Ascoli, G. A. 2005. Signal propagation in oblique dendrites of CA1 pyramidal cells. J Neurophysiol, 94, 4145–55.

Nevian, T., Larkum, M. E., Polsky, A. & Schiller, J. 2007. Properties of basal dendrites of layer 5 pyramidal neurons: a direct patch-clamp recording study. Nat Neurosci, 10, 206–14.

Palmer, L. M., Shai, A. S., Reeve, J. E., Anderson, H. L., Paulsen, O. & Larkum, M. E. 2014. NMDA spikes enhance action potential generation during sensory input. Nat Neurosci, 17, 383–90.

Ramaswamy, S. & Markram, H. 2015. Anatomy and physiology of the thick-tufted layer 5 pyramidal neuron. Front Cell Neurosci, 9, 233.

Ranganathan, G. N., Apostolides, P. F., Harnett, M. T., Xu, N. L., Druckmann, S. & Magee, J. C. 2018. Active dendritic integration and mixed neocortical network representations during an adaptive sensing behavior. Nat Neurosci, 21, 1583–1590.

Schaefer, A. T., Larkum, M. E., Sakmann, B. & Roth, A. 2003. Coincidence detection in pyramidal neurons is tuned by their dendritic branching pattern. J Neurophysiol, 89, 3143–54.

Schiller, J., Schiller, Y., Stuart, G. & Sakmann, B. 1997. Calcium action potentials restricted to distal apical dendrites of rat neocortical pyramidal neurons. Journal of Physiology-London, 505, 605–616.

Schiller, Y. 2002. Inter-ictal- and ictal-like epileptic discharges in the dendritic tree of neocortical pyramidal neurons. Journal of Neurophysiology, 88, 2954–2962.

Shai, A. S., Anastassiou, C. A., Larkum, M. E. & Koch, C. 2015. Physiology of Layer 5 Pyramidal Neurons in Mouse Primary Visual Cortex: Coincidence Detection through Bursting. Plos Computational Biology, 11.

Short, S. M., Oikonomou, K. D., Zhou, W. L., Acker, C. D., Popovic, M. A., Zecevic, D. & Antic, S. D. 2017. The stochastic nature of action potential backpropagation in apical tuft dendrites. J Neurophysiol, 118, 1394–1414.

Sjostrom, P. J. & Hausser, M. 2006. A cooperative switch determines the sign of synaptic plasticity in distal dendrites of neocortical pyramidal neurons. Neuron, 51, 227–38.

Sjostrom, P. J., Rancz, E. A., Roth, A. & Hausser, M. 2008. Dendritic excitability and synaptic plasticity. Physiol Rev, 88, 769–840.

Smith, S. L., Smith, I. T., Branco, T. & HÄUSSER, M. 2013. Dendritic spikes enhance stimulus selectivity in cortical neurons in vivo. Nature, 503, 115–120.

Spruston, N. 2008. Pyramidal neurons: dendritic structure and synaptic integration. Nat Rev Neurosci, 9, 206–21.

Stuart, G., Schiller, J. & Sakmann, B. 1997a. Action potential initiation and propagation in rat neocortical pyramidal neurons. J Physiol, 505 (Pt 3), 617–32.

Stuart, G., Spruston, N., Sakmann, B. & Hausser, M. 1997b. Action potential initiation and backpropagation in neurons of the mammalian CNS. Trends Neurosci, 20, 125–31.

Stuart, G. J. & Sakmann, B. 1994. Active Propagation of Somatic Action-Potentials into Neocortical Pyramidal Cell Dendrites. Nature, 367, 69–72.

Stuart, G. J. & Spruston, N. 2015. Dendritic integration: 60 years of progress. Nature Neuroscience, 18, 1713–1721.

Vetter, P., Roth, A. & Hausser, M. 2001. Propagation of action potentials in dendrites depends on dendritic morphology. J Neurophysiol, 85, 926–37.

Waters, J., Schaefer, A. & Sakmann, B. 2005. Backpropagating action potentials in neurones: measurement, mechanisms and potential functions. Prog Biophys Mol Biol, 87, 145–70.

Xu, N. L., Harnett, M. T., Williams, S. R., Huber, D., O’connor, D. H., Svoboda, K. & Magee, J. C. 2012. Nonlinear dendritic integration of sensory and motor input during an active sensing task. Nature, 492, 247–251.

Zhou, W. L., Short, S. M., Rich, M. T., Oikonomou, K. D., Singh, M. B., Sterjanaj, E. V. & Antic, S. D. 2015. Branch specific and spike-order specific action potential invasion in basal, oblique, and apical dendrites of cortical pyramidal neurons. Neurophotonics, 2.

